# GSK2556286 is a novel antitubercular drug candidate effective in vivo with the potential to shorten tuberculosis treatment

**DOI:** 10.1101/2022.02.04.479214

**Authors:** Eric L. Nuermberger, Maria Santos Martínez-Martínez, Olalla Sanz, Beatriz Urones, Jorge Esquivias, Heena Soni, Rokeya Tasneen, Sandeep Tyagi, Si-Yang Li, Paul J. Converse, Helena I. Boshoff, Gregory T Robertson, Gurdyal S Besra, Katherine A. Abrahams, Anna M Upton, Khisimuzi Mdluli, Gary W Boyle, Sam Turner, Nader Fotouhi, Nicholas C. Cammack, Juan Miguel Siles, Marta Alonso, Jaime Escribano, Joel Lelievre, Esther Pérez-Herrán, Robert H. Bates, Gareth Maher-Ewards, David Barros, Lluís Ballell, Elena Jiménez

## Abstract

As a result of a high-throughput compound screening campaign of *Mycobacterium tuberculosis* infected macrophages, a new preclinical drug candidate for the treatment of tuberculosis has been identified. GSK2556286 inhibits growth within human macrophages (IC_50_ = 0.07 µM), is active against extracellular bacteria in cholesterol-containing culture media and exhibits no cross-resistance with known antitubercular drugs. In addition, it has shown efficacy in different mouse models of tuberculosis (TB) and has an adequate safety profile in two preclinical species. These features indicate a compound with a novel mode of action, although still not fully defined, that is effective against both multidrug or extensively-resistant (M/XDR) and drug-sensitive (DS) *M. tuberculosis* with the potential to shorten the duration of treatment in novel combination drug regimens.

**One Sentence Summary:** GSK2556286 is a novel preclinical drug candidate for the treatment of tuberculosis with a new mode of action potentially able to contribute to the shortening of TB chemotherapy.

## Introduction

According to World Health Organization (WHO) estimates for 2019, 10 million people were newly diagnosed with tuberculosis (TB) and 1.4 million died (1), making TB the single greatest cause of death globally by a single infectious agent prior to the COVID-19 pandemic. Multidrug-resistant TB (MDR-TB) threatens TB control in many countries with approximately 363,000 new cases globally in 2019 (1). Furthermore, the incidence of extensively drug-resistant TB (XDR-TB), defined as MDR-TB plus resistance to at least one second-line injectable drug (e.g., amikacin, kanamycin or capreomycin) and a fluoroquinolone, was over 12,000 in 2019. Cases of XDR-TB have now been reported in over 100 countries (1). Despite regulatory approvals for bedaquiline (B), delamanid (D) and pretomanid (Pa) in the past decade to treat MDR or XDR-TB, there remains an unmet need for novel drugs with new mechanisms of action that are effective against drug-susceptible and drug-resistant forms of TB and shorten the duration of treatment required to prevent relapse.

To date, virtually all approved drugs used to treat TB were identified through phenotypic screens against actively replicating *Mycobacterium tuberculosis* in artificial nutrient-rich media, or they were repurposed from other infectious indications (2). The first-line TB drug pyrazinamide (Z) is the notable exception, having been identified by screening for activity in a murine TB model (3, 4). Few other pathogens rival *M. tuberculosis* in their ability to adapt to and persist within the infected host. Alternative screening methodologies that better represent the environmental conditions and stresses encountered by *M. tuberculosis* within the host have gained favor in recent years and may increase the efficiency with which new molecules with novel sterilizing activity are identified to complement existing TB drugs (5).

Over the last decade, we and others hypothesized that the macrophage, as a primary target of infection by *M. tuberculosis* and a niche in which the pathogen persists in established lesions, might represent an improved surrogate model to facilitate the discovery of novel TB drugs (6, 7). The cytochrome bc_1_:aa_3_ complex inhibitor telacebec is the first TB drug to reach clinical trials that was initially identified in a phenotypic high-content screening approach using a macrophage infection model (7, 8). Nonetheless, it is active against *M. tuberculosis* in standard nutrient-rich media as well as in macrophages. More recently, novel compounds with selective activity within macrophages were identified and shown to have cholesterol-dependent activity against extracellular *M. tuberculosis in vitro* (9). Previous observations suggest that cholesterol uptake and utilization is essential for pathogen survival in the host and indicate these pathways as potential targets for novel TB drugs (10, 11) Despite these encouraging results, no molecule identified as having such macrophage-specific, cholesterol-dependent activity *in vitro* has progressed to clinical proof-of-concept studies. Here, we describe the discovery of GSK2556286, a novel inhibitor of *M. tuberculosis* extracellularly in the presence of cholesterol and within human macrophages, that provides evidence of favorable *in vivo* efficacy and safety profiles justifying further development as an attractive companion drug with the potential to shorten the duration of treatment in novel combination regimens for drug-susceptible and drug-resistant TB.

## Results

### Microbiological profile

To identify compounds that effectively inhibit intracellular growth of *M. tuberculosis*, we screened a library of compounds against bacteria residing within human (THP-1) macrophage-like differentiated monocytes. The exploitation of this screening approach led to the identification of GSK2556286 (Fig. 1), a compound with potent activity (Table 1) against *M. tuberculosis* inside infected macrophages (lC_50_=0.07 µM in THP-l cells) and the unusual phenotype of requiring the presence of cholesterol to demonstrate activity in axenic culture (IC_50_=0.71-2.12 µM). The maximal % inhibition of growth achieved by GSK2556286 in these studies was 86% (range 62 to 89.4%).

**Table 1.** *In vitro* activity of GSK2556286 under various conditions

| Compound | $IC_{50}$ (μM) against *M. tuberculosis* strains H37Rv or Erdman | | | | |
| --- | --- | --- | --- | --- | --- |
|  | Intracellular activity | Extracellular activity |  |  |  |
|  | H37Rv in THP-1 cells | H37Rv in glucose media | H37Rv in cholesterol media | Erdman in glucose media | Erdman in cholesterol media |
| GSK2556286 | 0.07 | > 125 | 2.12 | > 50 | 0.71 |
| Rifampicin | 0.0008 | 0.15 | 0.2 | 0.1 | 0.1 |
| Moxifloxacin | 0.16 | 0.11 | 0.5 | 0.28 | 0.4 |

**Figure 1.**
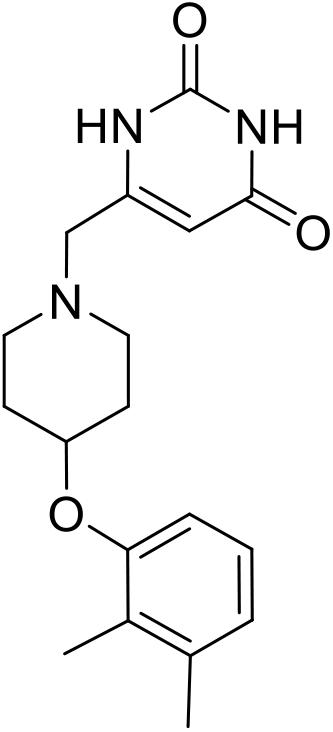
Chemical structure of GSK2556286

**Figure 2.**
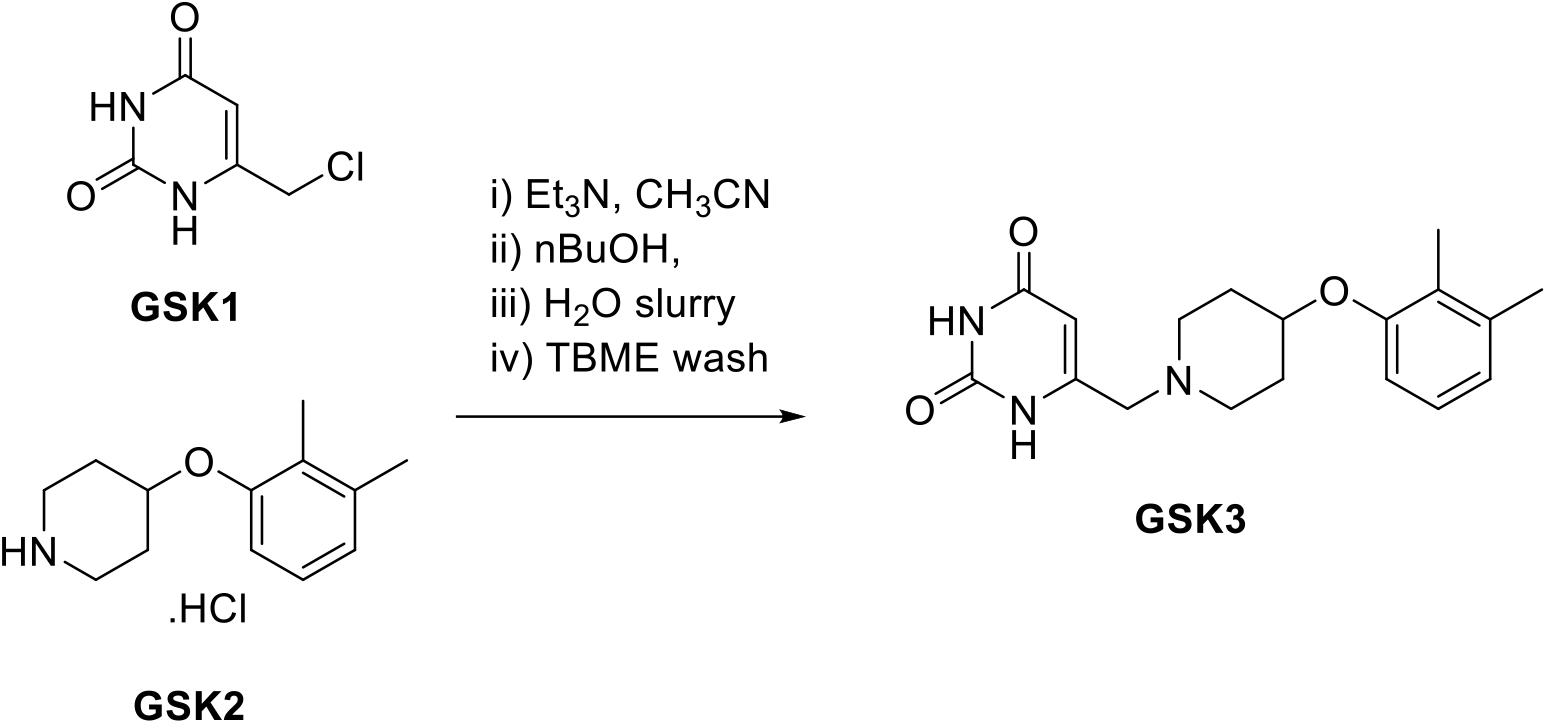
Synthetic route

GSK2556286 displayed consistent *in vitro* activity in the presence of cholesterol against a panel of clinical isolates with varying drug resistance phenotypes, including isolates from MDR and XDR-TB cases (Supplemental Table S1). The minimal inhibitory concentration of GSK2556286 that inhibited the growth of at least 90% of isolates (MIC90) was determined for 45 clinical isolates (from the National Institute of Health [NIH]), plus 3 laboratory strains, with different resistance phenotypes, including DS, MDR, XDR or other resistance phenotypes (Supplemental Table S2) as well as two additional species belonging to the *M. tuberculosis* complex, *Mycobacterium africanum (M. africanum)* and *Mycobacterium bovis (M. bovis)* in order to evaluate the activity of GSK2556286 on more genetically diverse species of the complex.

The MIC90 was 1.2 μM (MIC range 0.3 to 1.4 μM) similar to that determined for laboratory strains, Erdman and H37Rv (0.71 and 2.12 µM, respectively) in cholesterol containing media.

To investigate the potential mode of action, we isolated spontaneous resistant mutants to GSK2556286 when *M. tuberculosis* Erdman cultivated *in vitro* or extracted from the lungs of infected C3HeB/FeJ mice exposed to GSK2556286 at 96 µM (8xMIC in solid media including cholesterol). In total, 29 colonies isolated from GSK2556286-containing plates in the *in vitro* and *in vivo* experiments were serially passaged on GSK2556286-containing plates and confirmed to have IC_90_ values in the presence of cholesterol that were 10-fold higher than the wild-type parent. Whole genome sequencing and further analysis revealed that 14 out of 29 mutants had mutations mapping to the *Rv1625c* gene (*cya*) (Supplemental Table S3), which encodes a Class IIIa membrane-anchored adenylyl cyclase that is non-essential for growth under routine *in vitro* conditions and has been implicated in resistance to other compounds with cholesterol-dependent activity (9). The remainder of isolated resistant mutants remain under analysis to identify new mutations responsible of resistance.

None of the isolated resistant mutants, with *cya* mutation, had a complete deletion of the *cya* gene. Therefore, a *cya* knock-out mutant created in the H37Rv strain background was evaluated to confirm the role of *cya* in GSK2556286 resistance. The IC_50_ value in cholesterol media was >50 µM which is 25-fold higher than the IC_50_ value of the wild type strain. These results demonstrated that the *cya* gene has a role in resistance to GSK2556286 in *M. tuberculosis*.

Additional drug susceptibility testing of a selection of GSK2556286-resistant mutants (EM08, EM10, EM19, EM63) showed susceptibility to a selection of commonly used antitubercular drugs (Table 2, Supplemental Table S3) in axenic conditions but also showed susceptibility in an intra macrophage assay (Table 3).

**Table 2.**
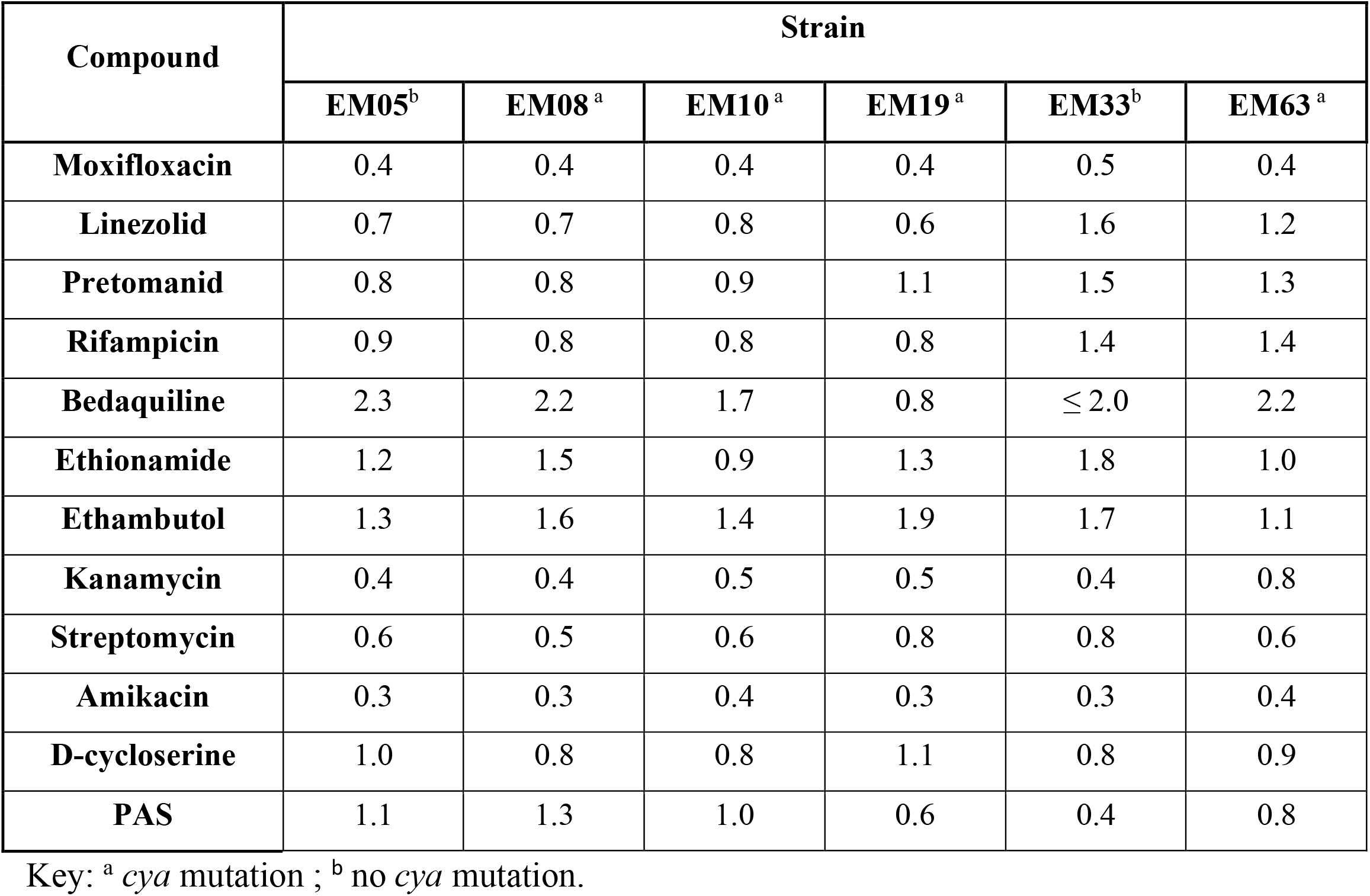
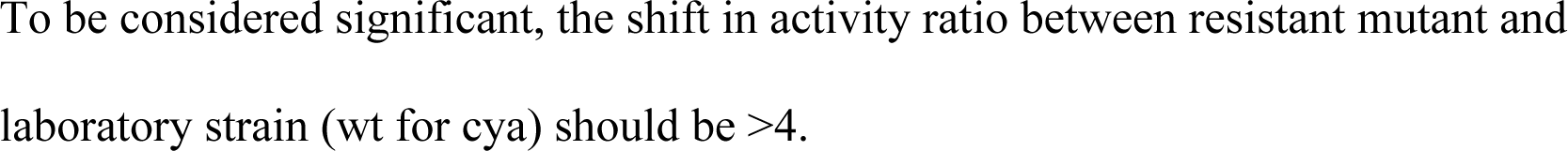
Antitubercular drug activity against selected GSK2556286-resistant *M. tuberculosis* strains. Data presented as the ratio of IC_90_ mutant/IC_90_ Wt EM01

**Table 3.** Susceptibility of THP-1 infected with a selection of GSK2556286-resistant *M. tuberculosis* strains to established antituberculars. Data presented as Ratio IC_50_ mutant/IC_50_ wild type EM01

|  | Strain |  |
| --- | --- | --- |
| Compound | EM19<sup>a</sup> | EM33<sup>b</sup> |
| Isoniazid | 0.57 | 0.54 |
| Rifampicin | 0.77 | 0.47 |
| Moxifloxacin | 0.32 | 0.30 |

Key: <sup>a</sup> cya mutation ; <sup>b</sup> no cya mutation.
To be considered significant, the shift in activity ratio between resistant mutant and laboratory strain (wt for cya) should be >4.

Calculation of the spontaneous frequency of resistance has, to date, been technically limited due to the challenging process of achieving acceptable growth in cholesterol-containing solid culture media. Further efforts are ongoing to refine the methodology to enable accurate assessments for spontaneous frequency of resistance to GSK2556286.

### Chemical and structural information and physicochemical properties

GSK2556286A (Fig. 1) is a substituted 4-aryloxypiperidine with a low-to-moderate molecular weight (MW=329.39).

The white-to-slightly-colored solid is crystalline with a high melting point (200°C) and a chromatographic logD and Pharmaceutical Formulation Index (PFI) of 4.4 and 6.4, respectively. GSK2556286A has excellent stability in the solid and solution states with respect to temperature and light, giving confidence that a solid oral product with a suitable shelf life can be developed. GSK2556286 is practically insoluble in water, fasted state simulated intestinal fluid (FaSSIF), fed state simulated intestinal fluid (FeSSIF) and aqueous solution in a pH range of 5-9. It is very slightly soluble in simulated gastric fluid and aqueous solution in a pH range of 2-4.

The Developability Classification System (DCS) class (12) borders on IIa/b at the predicted dose, suggesting potential issues with solubility and dissolution at high doses (Figure S1).

GSK2556286 was obtained in excellent yield in multi-gram scale trough a nucleophilic substitution of 6-(chloromethyl)-uracil derivative (GR202687X), and 4-(2,3-dimethylphenoxy)piperidine hydrochloride salt (GSK2422021A), both commercially available, catalyzed by triethylamine.

### In vitro absorption, distribution, metabolism and elimination (ADME) and pharmacokinetic (DMPK) profiles and potential for drug-drug interactions

Physicochemical properties, *in vitro* ADME, and *in vivo* DMPK profiles in preclinical species (mouse, rat, dog) were evaluated to support progression of GSK2556286 and dose prediction modelling in humans. GSK2556286 displayed notably higher solubility in simulated gastric fluid compared to that in other biologically relevant media. The compound exhibited high passive permeability in the hMDR1-MDCK-II cell line, and although it was shown to be an in vitro substrate for P-glycoprotein, based on its permeability and existing *in vivo* preclinical pharmacokinetic data, permeability is not expected to limit oral absorption of GSK2556286 in humans. Low intrinsic clearance (CL_int_) was determined in both human microsomes and hepatocytes. Low plasma protein binding (PPB) and low-to-moderate blood-to-plasma partitioning ratios (B/P) were observed in human and preclinical species (Supplemental Table S5).

Pharmacokinetics of GSK2556286 after intravenous and oral administration at various doses were evaluated in rodents and dogs. The compound exhibited low-to-moderate blood clearance, as predicted by CL_int_ in hepatocytes, and moderate volume of distribution. Absorption was rapid and oral bioavailability at pharmacologically relevant doses was high in mice and moderate in rats and dogs, in agreement with the expected first-pass effect (Supplemental Table S6)

Human PK parameters were calculated for GSK2556286 using a physiologically-based pharmacokinetic (PBPK) modelling approach (GastroPlus), based on physicopchemical, preclinical (in vitro and in vivo) and in vitro juman data. The PBPK models accurately predicted the IV and oral PK data from preclinical studies in mice, rats and dogs, and the prediction estimates a low human blood clearance (3.3 mL/min/kg), a moderate volume of distribution (3.5 L/kg) and high oral bioavailability (≥ 60% for predicted clinical doses). Taking as a reference the minimum AUC and C_max_ at 10 mg/kg associated with a maximum effect as a single drug in BALB/c mice, it was predicted a dose in humans between 150 and 300 mg/day administration (based on targeting AUC and C_max_, respectively).

To assess the risk of drug-drug interactions, direct inhibition of CYP isoforms was investigated by assessing the enzyme activities (CYP1A2, CYP2C9, CYP2C19, CYP2D6 and CYP3A4) in an incubation mixture of microsomes with NADPH, in the presence and absence of GSK2556286. This preliminary evaluation showed that, although it did not substantially inhibit CYP1A2, 2C9, 2C19, 2D6 and 3A4 (IC50 values >25 µM), there is a moderate risk of CYP3A4-mediated perpetrator drug interactions assuming a CYP3A4 IC50 value of 25 µM and predicted human PK parameters for a 150 mg dose of GSK2556286. (Supplemental Table S7).

All studies were conducted in accordance with the GSK Policy on the Care, Welfare and Treatment of Laboratory Animals and were reviewed the Institutional Animal Care and Use Committee either at GSK or by the ethical review process at the institution where the work was performed.

### Safety profile

GSK2556286 was evaluated in single dose oral toxicity studies in the rat, dog and cynomolgus monkey and in repeat dose oral toxicity studies of up to 4 weeks duration in the Wistar Han rat and cynomolgus monkey under GLP conditions and performed according to ICH guidelines. In addition, GSK2556286 was evaluated in a battery of *in vitro* and *in vivo* safety pharmacology (respiratory, cardiovascular and neurobehavioral tests) and genotoxicity studies (including an Ames test on GSK2422021, a synthetic intermediate, predicted degradant and a metabolite of GSK2556286).

In the definitive repeat dose oral toxicity studies in rat and monkey, adverse systemic effects were limited to the rats in the high dose group (1000 mg/kg/day). No adverse effects were observed at exposures up to an AUC_0-t_=65.2 µg*h/mL and C_max_=5.89 µg/mL in male and AUC_0-t_=129 µg*h/mL and C_max_=14.6 µg/mL in female rats and AUC_0-t_=158 µg*h/mL and mean C_max_=9.96 µg/mL in the cynomolgus monkey (gender-averaged). GSK2556286 did not produce acute cardiovascular effects in rat or monkey, respiratory effects in monkey or adverse neurobehavioural effects in rats in single or repeat dose studies up to 1000 mg/kg/day and the weight of evidence from *in vitro* and *in vivo* assessments indicates that GSK2556286 does not present a genotoxic hazard to humans. The preclinical safety profile supports continued progression to a first-time-in-humans (FTIH) trial.

All animal studies were ethically reviewed and carried out in accordance with Animals (Scientific Procedures) Act 1986 and the GSK Policy on the Care, Welfare and Treatment of Animals.

### Efficacy in murine models of TB

#### Dose-ranging activity as monotherapy

GSK2556286 was tested for *in vivo* efficacy in two different murine models of chronic TB infection. In BALB/c mice, *M. tuberculosis* infection promotes development of inflammatory cellular lung lesions in which *M. tuberculosis* resides virtually entirely intracellularly, especially in macrophages, including foamy macrophages. In contrast, C3HeB/FeJ mice also form caseating granulomatous lung lesions, in which *M. tuberculosis* is found extracellularly in the acellular central caseum as well as inside neutrophils and macrophages (13, 14).

The compound showed a statistically significant bactericidal effect when used as a single agent for 1 month (4 weeks) in chronic infection models in both mouse strains (Table 4). In BALB/c mice, all GSK2556286 doses tested were superior to no treatment (p< 0.0001) and the bactericidal effect was similar in magnitude to that of isoniazid. The maximal effect was achieved at a dose less than or equal to 10 mg/kg, which corresponded to a C_max_ of 1.38 ug/mL and an AUC_last_ of 6.61 µg*h/mL. In C3HeB/FeJ mice, all doses above 10 mg/kg were significantly better than no treatment (p<0.05) before adjustment for multiple comparisons, although only the 40 mg/kg dose was significantly different from no treatment after adjusting for multiple comparisons. Isoniazid was superior to each dose of GSK2556286 in C3HeB/FeJ mice (p<0.01).

**Table 4.** Lung CFU counts in BALB/c and C3HeB/FeJ mice after 4 weeks of GSK2556286 treatment

| | Mean (± SD) lung CFU counts | | | | | |
| --- | --- | --- | --- | --- | --- | --- |
|  | BALB/c mice |  |  | C3HeB/FeJ mice |  |  |
| Regimen | Week -8 | Day 0 | Month 1 | Week -8 | Day 0 | Month 1 |
| Untreated | 1.95 ± 0.14 | 6.84 ± 0.23 | 6.71 ± 0.34 | 1.68 ± 0.10 | 7.00 ± 0.77 | 7.42 ± 0.95 |
| INH (10 mg/kg) | | | 5.54 ± 0.41* | | | 4.93 ± 0.46* |
| GSK'286 (10 mg/kg) | | | 5.63 ± 0.12* | | | 6.52 ± 0.93 |
| GSK'286 (40 mg/kg) | | | 5.69 ± 0.05* | | | 6.30 ± 0.54<sup>#</sup> |
| GSK'286 (100 mg/kg) | | | 5.82 ± 0.08* | | | 6.31 ± 1.00 |
| GSK'286 (200 mg/kg) | | | 5.50 ± 0.18* | | | 6.30 ± 0.96 |

Key: \* $p < 0.01$ compared to untreated group. <sup>#</sup> $p < 0.05$ compared to untreated group
GSK'286=GSK2556286

### Contribution to bactericidal activity in combination therapy

Given the requirement for combination chemotherapy in the treatment of TB and the urgent need for novel regimens comprised of drugs that retain activity against MDR- and XDR-TB strains, the efficacy of GSK2556286 was evaluated in a subacute infection model in BALB/c mice that enables the evaluation of drug regimens against a higher bacterial burden (15). GSK2556286 (50 mg/kg) was co-administered with bedaquiline (B) and pretomanid (Pa) and the efficacy of this regimen was compared to that of BPa plus linezolid (L), which comprises a novel short-course regimen (16, 17) that was recently approved for treatment of XDR-TB and refractory MDR-TB. The addition of GSK2556286 to the BPa combination significantly increased efficacy, compared to BPa alone, after two months of treatment (p<0.001) (Table 5).

**Table 5.**
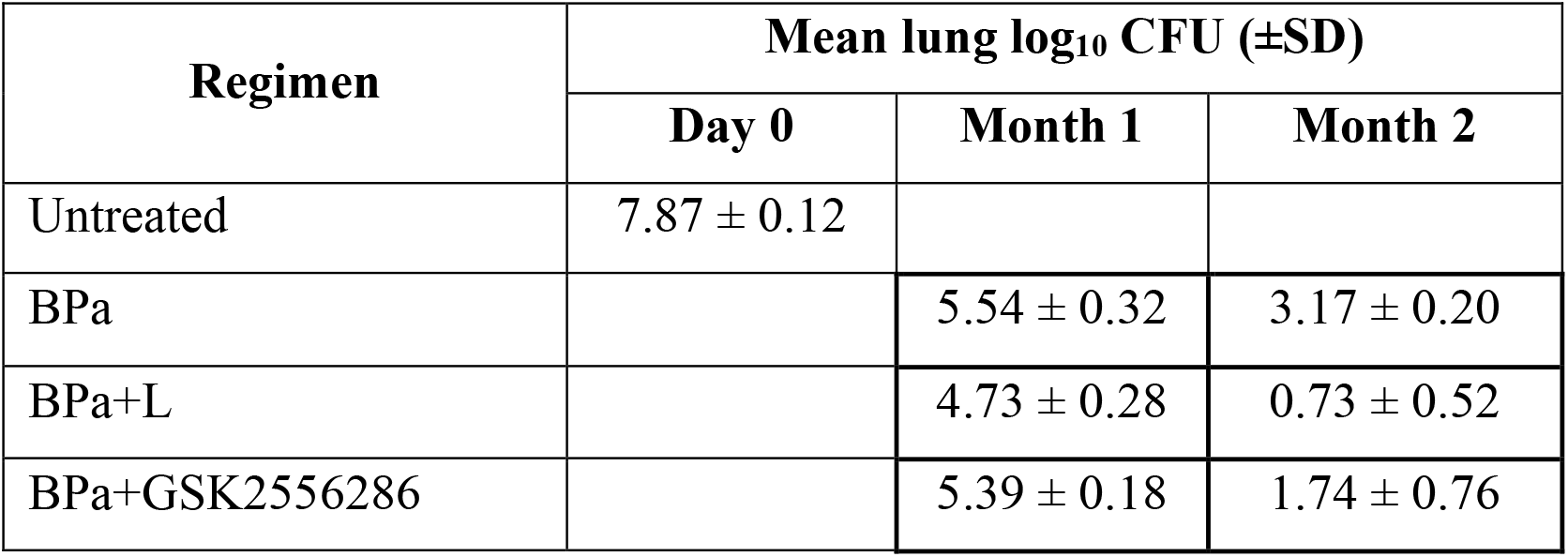
Efficacy of GSK2556286 when combined with B and Pa in a BALB/c mouse model of TB. For comparison, the three-drug combination BPa+L is included

### Contribution to treatment-shortening activity in combination therapy

Although the bactericidal activity of this novel 3-drug combination was not as great as that of BPaL (p<0.01), the 6-log_10_ magnitude of the killing effect and clear contribution of GSK2556286 to the combination led us to assess the potential of GSK2556286 to contribute sterilizing activity when incorporated into 3- and 4-drug regimens with B, Pa and L in the subacute BALB/c mouse infection model, using the proportion of mice with relapse-free cure as the primary endpoint. The standard of care RHZ (rifampicin+isoniazid+pyrazinamide) was also included as reference for bactericidal outcome after 1 and 2 months of treatment and relapse endpoint after 4 months based on previous data indicating RHZ requires more than 3 months to observe significant reductions in the proportion of mice that relapse (18, 19).

After 2 months of treatment, regimens combining GSK2556286 with BPa, BL or BPaL resulted in significantly lower lung CFU counts compared to the first-line RHZ control (p<0.0001, p=0.0417, and p<0.0001, respectively) (Table 6). BPa+GSK2556286 and BPaL+GSK2556286 were not significantly different from BPaL at this time point, but BL+GSK2556286 and PaL+GSK2556286 were significantly less active than BPaL (p<0.0001). With respect to the relapse outcome, treatment with BPaL, BPa+GSK2556286 and BPaL+GSK2556286 for 2 months resulted in lower proportions of mice relapsing compared to treatment with RHZ for 4 months, indicating the treatment-shortening potential of regimens combining GSK2556286 with BPa and BPaL, compared to RHZ. BL+GSK2556286 required 3 months of treatment to achieve a relapse rate lower than RHZ for 4 months.

**Table 6.**
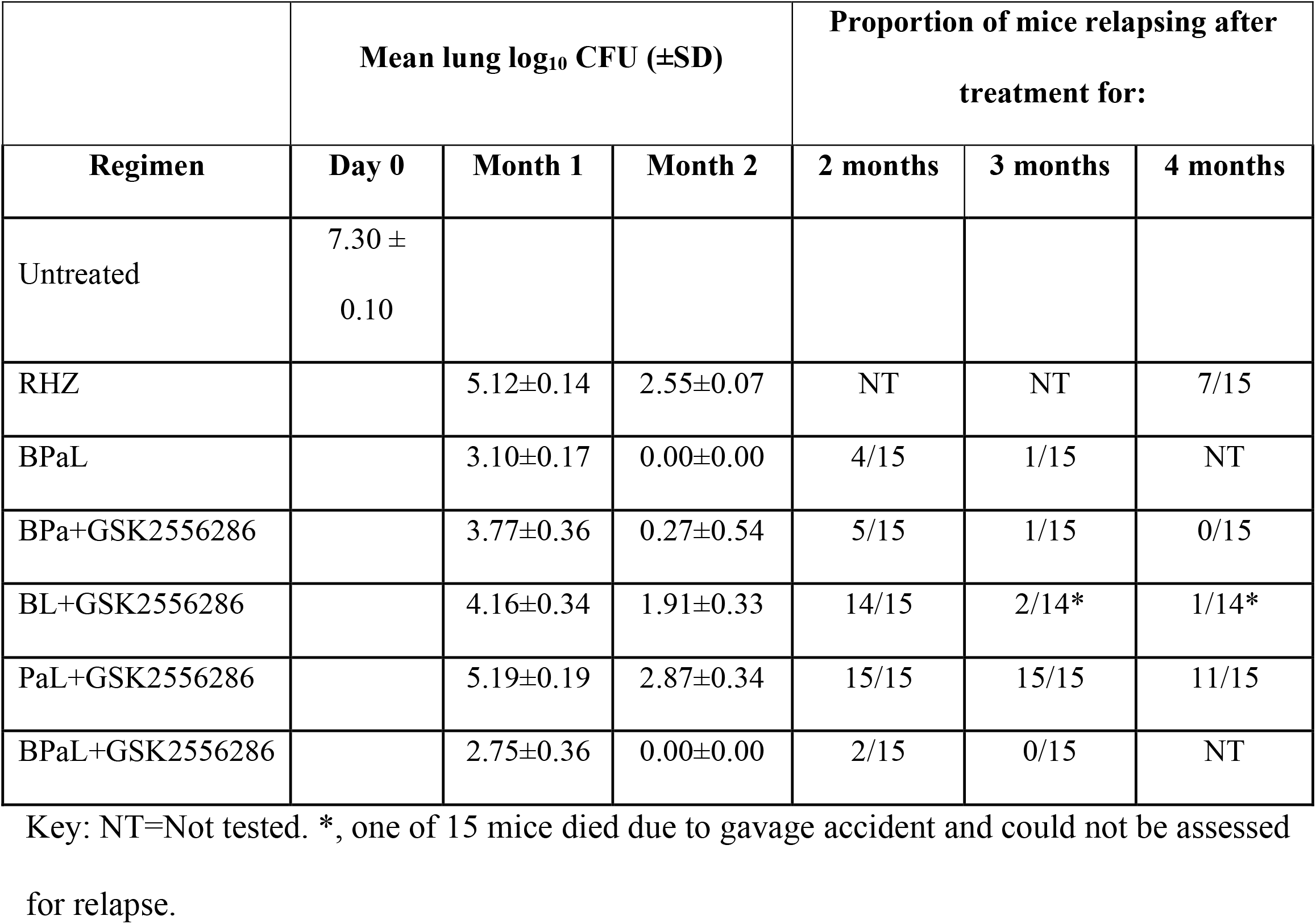
Efficacy of GSK2556286 when combined with various 2- and 3-drug combinations of B, Pa and L in a BALB/c mouse model of TB.

| Regimen | Mean lung log<sub>10</sub> CFU (±SD) |  |  | Proportion of mice relapsing after treatment for: |  |  |
| --- | --- | --- | --- | --- | --- | --- |
|  | Day 0 | Month 1 | Month 2 | 2 months | 3 months | 4 months |
| Untreated | 7.30 ± 0.10 |  |  |  |  |  |
| RHZ |  | 5.12±0.14 | 2.55±0.07 | NT | NT | 7/15 |
| BPaL |  | 3.10±0.17 | 0.00±0.00 | 4/15 | 1/15 | NT |
| BPa+GSK2556286 |  | 3.77±0.36 | 0.27±0.54 | 5/15 | 1/15 | 0/15 |
| BL+GSK2556286 |  | 4.16±0.34 | 1.91±0.33 | 14/15 | 2/14* | 1/14* |
| PaL+GSK2556286 |  | 5.19±0.19 | 2.87±0.34 | 15/15 | 15/15 | 11/15 |
| BPaL+GSK2556286 |  | 2.75±0.36 | 0.00±0.00 | 2/15 | 0/15 | NT |

Key: NT=Not tested. \*, one of 15 mice died due to gavage accident and could not be assessed for relapse.

The proportions of mice relapsing after 2 and 3 months of treatment with BPaL, BPa+GSK2556286 and BPaL+GSK2556286 did not significantly differ, indicating that GSK2556286 could replace L in the BPaL regimen without loss of efficacy. On the other hand, PaL+GSK2556286 was associated with significantly more relapses (p<0.0001) at all time points and BL+GSK2556286 was associated with more relapses (p=0.0005) after 2 months of treatment but was similar after 3 and 4 months.

### Activity in combination therapy in C3HeB/FeJ mouse model

These 3- and 4-drug regimens in which GSK2556286 was either added to BPaL or replaced B, Pa or L were also evaluated for CFU reduction in C3HeB/FeJ mice infected with *M. tuberculosis* H37Rv. After one month of treatment, none of these regimens was statistically significantly different from the RHZ or BPaL regimens (Figure 3). After two months, only PaL+GSK2556286 was significantly worse than RHZ (p=0.0006), suggesting GSK2556286 could replace either Pa or L in the BPaL regimen without loss of efficacy compared to BPaL or RHZ (group mean CFU counts are presented in Supplemental).

**Figure 3.**
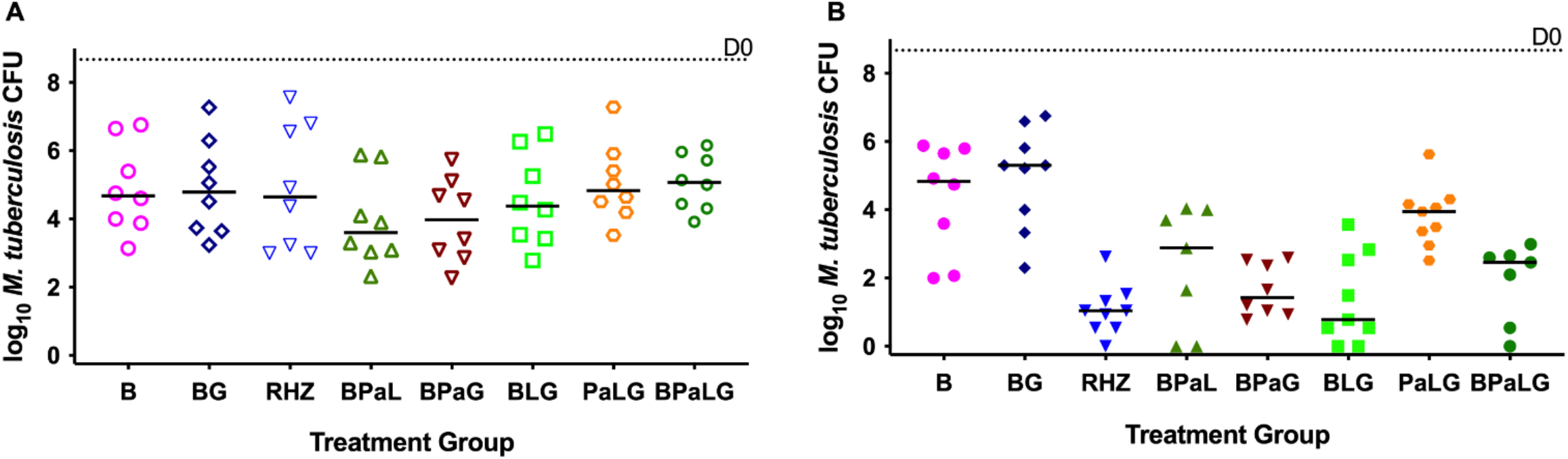
Efficacy of GSK2556286 (G) when combined with the various 2- and 3-drug combinations of B (bedaquiline), Pa (pretomanid) and L (linezolid) for 1 month (A) or 2 months (B) in a C3HeB/FeJ mouse model of TB. For comparison, B, BG and BPaL are included. Bars indicates median CFU counts. Median CFU on Day 0, the start of treatment, is indicated by dotted line.

## Discussion

GSK2556286 is a novel, small molecule, antitubercular compound identified from high-throughput intramacrophage screening that is active against a variety of drug-sensitive and drug-resistant clinical isolates in axenic culture in the presence of cholesterol as carbon source. Cholesterol uptake, catabolism and broader utilization are important for maintenance of the pathogen in the host, and other inhibitors of *M. tuberculosis* with activity revealed in the presence of cholesterol have been identified (9, 10, 11).

To further understand the potential cholesterol-dependent mode of action, GSK2556286-resistant *M. tuberculosis* clones were isolated on media containing cholesterol as the primary carbon source and analyzed by whole genome sequencing. Approximately half of the resistant clones sequenced harbored mutations in the gene for the membrane-anchored adenylyl cyclase, *cya*, without altering the viability of the bacteria at least in laboratory conditions. The observed frameshift and premature stop codon mutations indicated that loss of *cya* function results in resistance to GSK2556286 and this was confirmed with the *cya* knockout-mutant. These findings are similar to other recently discovered cholesterol-dependent antitubercular leads that directly activate *cya* and induce bacterial cAMP production and inhibit cholesterol production in wild type *M. tuberculosis* in an *cya*-dependent fashion [9]. Increased intracellular levels of cAMP, one of the main secondary messengers in the cell, may negatively regulate cholesterol and propionate utilization by *M. tuberculosis*, reducing bacterial growth when it is dependent upon utilizing these carbon sources (9, 20, 21). Therefore, GSK2556286 may likewise act by activating *cya*, inducing cAMP production and negatively regulating cholesterol and propionate utilization. Ongoing studies to further evaluate the mode of action of GSK2556286, including its effects on cAMP levels and its impact in the presence of cholesterol will be reported separately. Despite the need for further elucidation of the specific mechanism of action, GSK2556286-resistant mutants remained susceptible to a list of well-known antitubercular drugs that suggests the novelty of this mechanism.

Given that the macrophage is a major cellular niche for *M. tuberculosis*, blocking replication of the bacterium in this environment may enhance existing or future TB drug regimens (2, 6). However, human TB disease is also characterized by development of caseation necrosis leading to closed caseous foci and cavities in which *M. tuberculosis* is found extracellularly, in caseum. As caseum is also rich in cholesterol, those bacilli persisting extracellularly in the acellular zones of caseous foci could also be susceptible to GSK2556286, as they are in axenic culture in the presence of cholesterol (22, 23).

To demonstrate this therapeutic potential, GSK2556286 was evaluated, alone and in combination with other drugs, in two murine TB models, one of which (C3HeB/FeJ mice) is distinguished from other mouse models by its propensity to develop caseating lung lesions (24-26). When used alone, GSK2556286 exhibited bactericidal effects in chronic infection models in both BALB/c mice, where virtually all bacteria reside intracellularly, and in C3HeB/FeJ mice, which form large caseating granulomas in which most bacteria are extracellular in caseum.

When used in combination to identify novel efficacious combination drug regimens including GSK2556286 in both models of TB, BALB/c and C3HeB/FeJ mice, GSK2556286 exhibited its potential to replace linezolid (L) in the BPaL regimen without loss of sterilizing activity. BPaL has been recently approved as a novel short-course oral treatment for XDR-TB and treatment refractory MDR-TB (27).

Although the individual contribution of GSK2556286 to the regimen’s sterilizing activity was not shown directly in these studies, there was no loss of bactericidal or sterilizing activity of the BPaL regimen when GSK2556286 was substituted for linezolid in the BALB/c mouse infection model. The contribution of linezolid to the sterilizing activity of the BPaL regimen has been repeatedly demonstrated in this model (17) and thus these data suggest that GSK2556286 is likely to be contributing to the overall efficacy of the BPa+GSK2556286 regimen.

Furthermore, although there were more relapses after 2 and 3 months of treatment when GSK2556286 was substituted for Pa in BPaL (14 vs 4, and 2 vs 1, respectively), BL+GSK2556286 treatment for 3 and 4 months resulted in fewer relapses (2 and 1, respectively) than RHZ treatment for 4 months (7 relapses) in BALB/c mice and BL+GSK2556286 had bactericidal activity that could not be distinguished from RHZ or BPaL in C3HeB/FeJ mice.

In summary, GSK2556286 acts via a novel model of action to achieve significant in vivo activity in murine models displaying both cellular and extracellular lesion compartments. This result combined with the compound’s low clearance values across a number of species, low propensity towards drug-drug interaction liabilities and adequate preliminary toxicology profile (genotoxicity, safety pharmacology, general toxicology) present evidence supporting its progression as a new clinical candidate for the treatment of both MDR and drug-susceptible TB that has potential to contribute to the shortening of TB chemotherapy.

## Materials and Methods

### Microbiological assays

The MIC of GSK2556286 against extracellular *M. tuberculosis* laboratory strains was determined in standard media with glucose as a carbon source and also in media supplemented with cholesterol. For MIC determination in the absence of cholesterol, approximately 1×10^5^ CFU/mL of *M. tuberculosis* H37Rv (ATCC 25618) or Erdman (TMCC 107) was added to 96-well flat-bottom plates containing ten two-fold drug dilutions of GSK2556286 in Middlebrook 7H9 medium (Difco) supplemented with 2% glucose, 0.025% Tween 80, 0.05% Tyloxapol and 10% albumin-dextrose-catalase [ADC]). Plates were placed in a sealed box to prevent drying and incubated at 37°C for 6 days. 25 µL of Resazurin solution (38.6 µM) (Resazurin Tablets for Milk Testing; Ref 330884Y’ VWR International Ltd, in 30 mL of phosphate buffered saline [PBS]) were added to each well. Fluorescence was measured after 48 hours at 37ºC using a SpectraMax M5 microplate reader (Molecular Devices). Non-linear regression analysis was used to fit the normalized fluorescence results into dose response curves and IC_50_ and IC_90_ values were determined using Excel Add-in XLFit. Reported data are the average of at least two experiments.

The MIC of GSK2556286 in media containing cholesterol as the carbon source was determined against *M. tuberculosis* H37Rv and Erdman strains with a final inoculum of approximately 1.4 ×10^6^ CFU/mL. Bacteria grown in 7H9 medium supplemented with 2% glucose and 0.025% Tyloxapol to an optical density (OD) around 0.5 at 600 nm were pelleted by centrifugation and washed twice with cholesterol media (Middlebrook 7H9 supplemented with 1 g/L KH_2_PO_4_, 2.5 g/L Na_2_HPO_4_, 0.5 g/L asparagine, 50 mg/L ferric ammonium citrate, 10 mg/L MgSO_4_·7H_2_O, 0.5 mg/L CaCl_2_, and 0.1 mg/L ZnSO_4_ and 0.01% cholesterol as the sole carbon source). Pellets were resuspended in cholesterol media and incubated for at least 3 days at 37ºC. Plates containing GSK2556286 were inoculated and incubated at 37ºC for an additional 7 days, resazurin solution (38.6 µM) was added and incubated for 48 hours before fluorescence was measured. Rifampicin (Sigma R3501) was used as positive control up to 0.36 µM. IC_50_ and IC_90_ values were determined as described above. Reported data are the average of at least two experiments.

The intracellular antibacterial activity of GSK2556286 was determined using human THP-1 cells maintained in RPMI-1640 media containing 10% fetal bovine serum (FBS), 1mM of pyruvate, and 2mM of L-glutamine, and incubated at 37 ºC with 5% CO_2_. An *M. tuberculosis* H37Rv reporter strain carrying the firefly luciferase gene (under the control of the *hsp60* promoter) was grown in Middlebrook 7H9 broth supplemented with 10% ADC, 0.4% glycerol and 0.05% Tween 80 until the mid-log phase. THP-1 cells were infected at a multiplicity of infection (MOI) of 1:1 in antibiotic-free RPMI-1640 media containing 10% FBS, 1 mM of pyruvate, 2 mM of L-glutamine and 20 nM of phorbol 12-myristate 13-acetate (PMA) for 4 hours at 37ºC with 5% CO_2_. Following a 4-hour incubation period, infected cells were harvested and plated onto 96-well plates containing either GSK2556286 (up to 25 µM), rifampicin up to 0.73 µM as positive control, or dimethyl sulfoxide (DMSO) (<0.5% final concentration). After 5 days of incubation, cell luminescence was measured using the Bright-Glo Promega kit and SpectraMax M5 plate reader. The percentages of inhibition were calculated relative to the DMSO control well. For each compound, the average value of the duplicate samples was calculated, and a sigmoidal dose-response (variable slope) curve was fit by nonlinear regression (GraphPad) to enable estimation of the IC_50_.

### MIC determination against a panel of clinical isolates on cholesterol-based media

The MIC of GSK2556286 was determined against 45 clinical isolates of *M. tuberculosis* with different resistance phenotypes maintained by the National Institutes of Health. *M. tuberculosis* strains HN878, CDC1551, Erdman and H37Rv were included as controls. Individual *M. africanum* and *M. bovis* isolates were included to represent other members of the *M. tuberculosis* complex. Briefly, bacteria were grown to an OD of 0.2-0.6 in the 7H9 medium supplemented with bovine serum albumin, Tyoxapol and cholesterol as a sole carbon source and added (2 × 10^4^ bacteria per well) to 96-well plates containing GSK2556286 (in concentrations ranging up to 50 µM). Para-aminosalicylic acid (PAS), isoniazid, and bedaquiline were used as positive controls and DMSO as a negative control. Plates were incubated for up to 3 weeks at 37°C. At various time points, plates were read with an inverted enlarging mirror plate reader and graded as either growth or no growth to determine the MIC. The time point was dependent on the growth rate of the strain in the drug-free control medium (generally between 1-2 weeks). After 2 weeks of incubation, resazurin was added to plates. Following incubation at 37°C for 24 hours, results were read visually with an inverted enlarging mirror plate reader (blue = growth inhibition, pink= growth). The lowest concentration to inhibit growth was defined as the MIC.

### Selection of GSK2556286 resistant mutants in vitro

The MIC of GSK2556286 in cholesterol-containing agar medium was used to establish the concentration of GSK2556286 for the selection of resistant mutants. Stocks of *M. tuberculosis* Erdman (2×10^9^ CFU/mL) were thawed and diluted 1:5 in PBS, and 0.1 mL aliquots were plated on 7H11 agar media containing cholesterol or cholesterol plus dextrose as carbon sources, with or without GSK2556286 at 96 µM (8xMIC). The bacterial colonies were counted after 4 or 6 weeks of incubation.

For genetic characterization, genomic DNA from single colonies was extracted with the MasterPureTM DNA Purification Kit from Epicentre (Cat. No. MCD85201) according to the manufacturer’s instructions. DNA libraries were generated following the Nextera XT Illumina protocol (Nextera XT Library Prep kit (FC-131-1024)). 0.2ng/ul purified gDNA was used to initiate the protocol. The multiplexing step was performed using Nextera XT Index Kit (FC-131-1096). The libraries were sequenced using a 2×150pb paired-end run, NextSeq high output reagent kit on a NextSeq Sequencer according to manufacturer’s instructions (Illumina). Quality assessment was performed using prinseq-lite program (28) applying following parameters: Min_length: 50, Trim_qual_right: 20, Trim_qual_type: mean and Trim_qual_window: 20. R1 and R2 from Illumina sequencing where joined using fastq-join from ea-tools suite (29).

### Creation of Rv1625c knock-out mutant

A knock-out mutant was created in the H37Rv strain by replacing *Rv1625c* with a hygromycin resistance cassette using the recombineering approach developed by Murphy et al (30). Gene replacement was confirmed by PCR.

### Selection of GSK2556286-resistant mutants in vivo

C3HeB/FeJ mice received a low-dose aerosol infection with 31 CFU/ lung of *M. tuberculosis* Erdman. Starting 8 weeks post-infection, mice were given oral doses of GSK2556286 at 100 mg/kg, 5 days a week for 6 weeks. Lungs were harvested and homogenates were prepared and plated in serial 10-fold dilutions on plates with and without GSK2556286 at 100 µM.

### Mouse efficacy studies

All housing and procedures involving mice were approved by the Institutional Animal Care and Use Committee at Johns Hopkins University School of Medicine and the GSK Policy on the Care, Welfare and Treatment of Animals.

### Mice

Female specific pathogen-free BALB/c mice and C3HeB/FeJ mice, each aged 5-6 weeks, were purchased from Charles River (Wilmington, MA) and Jackson Laboratories (Bar Harbor, ME), respectively. Mice were housed in a bio-safety level 3 animal facility.

### Mycobacterial Strain. M. tuberculosis

H37Rv was mouse-passaged, frozen in aliquots and sub-cultured in Middlebrook 7H9 broth with 10% oleic acid-albumin-dextrose-catalase (OADC) (Fisher, Pittsburgh, PA) and 0.05% Tween 80 prior to infection.

### Infection

BALB/c and C3HeB/FeJ mice were infected with *M. tuberculosis* H37Rv, using the Inhalation Exposure System (Glas-col, Terre Haute, IN). The chronic infection model in both mouse strains was initiated with a thawed aliquot of the bacterial culture that was diluted 1:50 with sterile PBS for infecting BALB/c mice and 1:100 for infecting C3HeB/FeJ mice with the goal of implanting approximately 100 and 50-75 CFU, respectively, in the lungs. Mice were held for 6 weeks before beginning treatment. The subacute infection model in BALB/c mice was initiated with a late log-phase culture in 7H9 broth (optical density at 600 nm of 0.8-1) with the goal of implanting 3.5-4 log_10_ CFU. Mice were held for 2 weeks before beginning treatment.

### Drug treatments

For dose-ranging monotherapy studies, GSK2556286 in doses of 10, 40, 100 and 200 mg/kg body weight was formulated in 1% methylcellulose and isoniazid 10 mg/kg was prepared in distilled water. Drugs were administered orally via gavage once daily, five days per week, for four weeks. Positive and negative controls received isoniazid and no treatment, respectively.

For experiments evaluating the activity of GSK2556286 in combinations with bedaquiline, pretomanid and linezolid, GSK2556286 and isoniazid were formulated as described above. Other drugs were formulated and administered at the indicated dose as previously described (16, 17): rifampicin (10 mg/kg) and pyrazinamide (150 mg/kg) in distilled water, bedaquiline (25 mg/kg) in acidified 10% (2-Hydroxypropyl)-ß-cyclodextrin (HPCD) solution, pretomanid (100 mg/kg) in 20% HPCD and lecithin (CM-2) formulation and linezolid (100 mg/kg) in 0.5% methylcellulose.

### Assessment of efficacy

Two microbiological outcomes were assessed: lung CFU counts during treatment and the proportion of mice relapsing after completion of treatment. Lungs were collected and homogenized in glass grinders at pre-specified time points during and after drug treatment. The homogenates were serially diluted in PBS and plated on Middlebrook 7H11 agar plates supplemented with 10% (v/v) OADC (GIBCO) and cycloheximide [10 mg/mL], carbenicillin [50 mg/mL], polymixin B [25 mg/mL] and trimethoprim [20 mg/mL]. Homogenates from mice receiving drug combinations were plated on the same agar media but with the addition of activated charcoal powder (0.4%w/v) to prevent carryover of bedaquiline. Colonies were counted after 4 and 6 weeks of incubation at 37°C to ensure all cultivable bacteria would be detected. Relapse after 2, 3 and 4 months of treatment with drug combinations was assessed by holding cohorts of 15 mice per group for an additional 3 months without treatment before sacrificing the mice and plating the entire lung homogenate, as described above. Relapse was defined by detection of ≥ 1 CFU.

### Statistical analysis

CFU counts (x) were log-transformed (as x+1) before analysis. Group means were compared by one-way analysis of variance with Dunnett’s post-test to control for multiple comparisons. Group relapse proportions were compared using Fisher’s exact test, adjusting for multiple comparisons. The Kruskal-Wallis test and Dunn’s non-parametric post-test to adjust for multiple comparisons were used to test for significance on non-normally distributed CFU data from C3HeB/FeJ mice. GraphPad Prism version 6 (GraphPad, San Diego, CA) was used for all analyses. Use of 15 mice per group for relapse assessment provides approximately 80% power to detect 40 percentage point differences in the relapse rate, after setting alpha at 0.01 to adjust for up to 5 simultaneous two-sided comparisons. Smaller differences may not be meaningful in terms of shortening the duration of treatment.

Data will be made publicly available upon publication and upon request for peer review.

## Acknowledgments

We would like to thank to Kevin Pethe and Jichan Jang for their collaboration with the screening at Institute Pasteur Korea. We would like to thank Ken Duncan and Peter Warner from the Bill & Melinda Gates Foundation, Steve Berthel from the New Venture Fund, and GSK technical and administrative support staff. We would like to thank Dirk Schanppinger and Curtiss Engelhart for providing KO strain and Anne Lenaerts for assistance with the in vitro and *in vivo* selection of resistant mutants.

